## Supplementary Figure 1 for "Embedding stochastic dynamics of the environment in spontaneous activity by prediction-based plasticity"

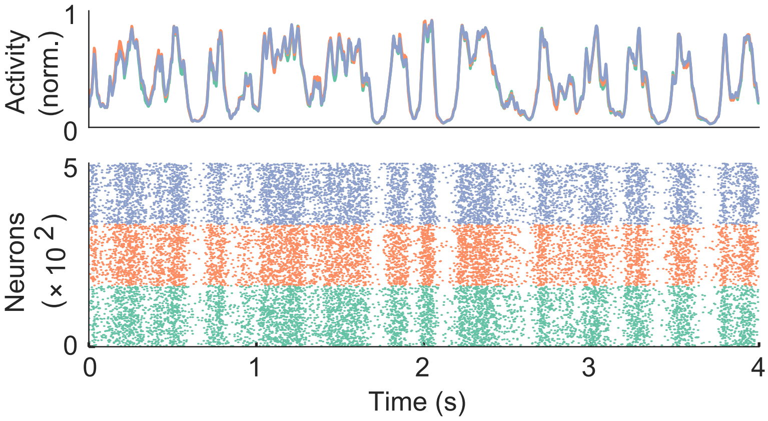


**Supplementary Figure 1. Unstructured spontaneous activity before learning.** Example spontaneous assembly reactivations (top) and raster plot (bottom) of the network are shown. Colors indicate the corresponding stimulus patterns shown in Fig.2a.
