## Supplementary Figure 2 for "Embedding stochastic dynamics of the environment in spontaneous activity by prediction-based plasticity"

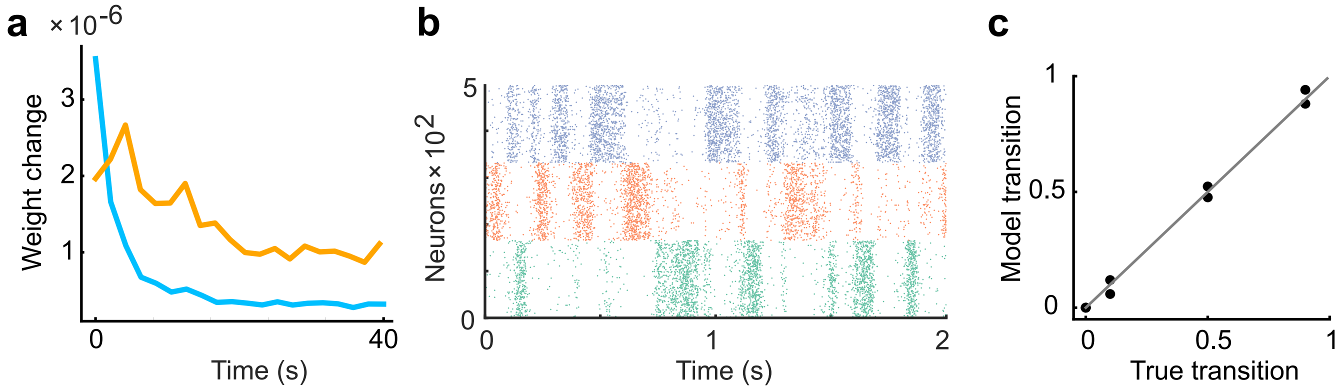


**Supplementary Figure 2. Inhibitory plasticity lags behind excitatory plasticity.** (a) The learning rate of inhibitory plasticity was made twice that of excitatory plasticity.The inhibitory plasticity still occurred on a slower timescale than excitatory plasticity (Froemke, 2007). (b) Example raster plot of spontaneous assembly reactivations of the learned network are shown. (c) Even if the learning rate of inhibitory plasticity was larger, the spontaneous activity reproduced transition statistics of external stimulus patterns.
