## Supplementary Figure 3 for "Embedding stochastic dynamics of the environment in spontaneous activity by prediction-based plasticity"

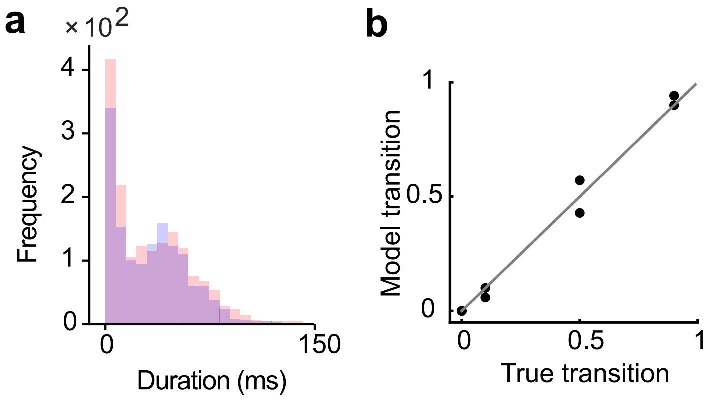


**Supplementary Figure 3. The network performance is less sensitive to the duration of evoked assembly activations during learning.** (a) The distributions of the durations of the assembly reactivations after training with input states of half the duration of the initial setting are shown. (b) The spontaneous activity of the trained network still showed an appropriate transition dynamics.
