## Supplementary Figure 4 for "Embedding stochastic dynamics of the environment in spontaneous activity by prediction-based plasticity"

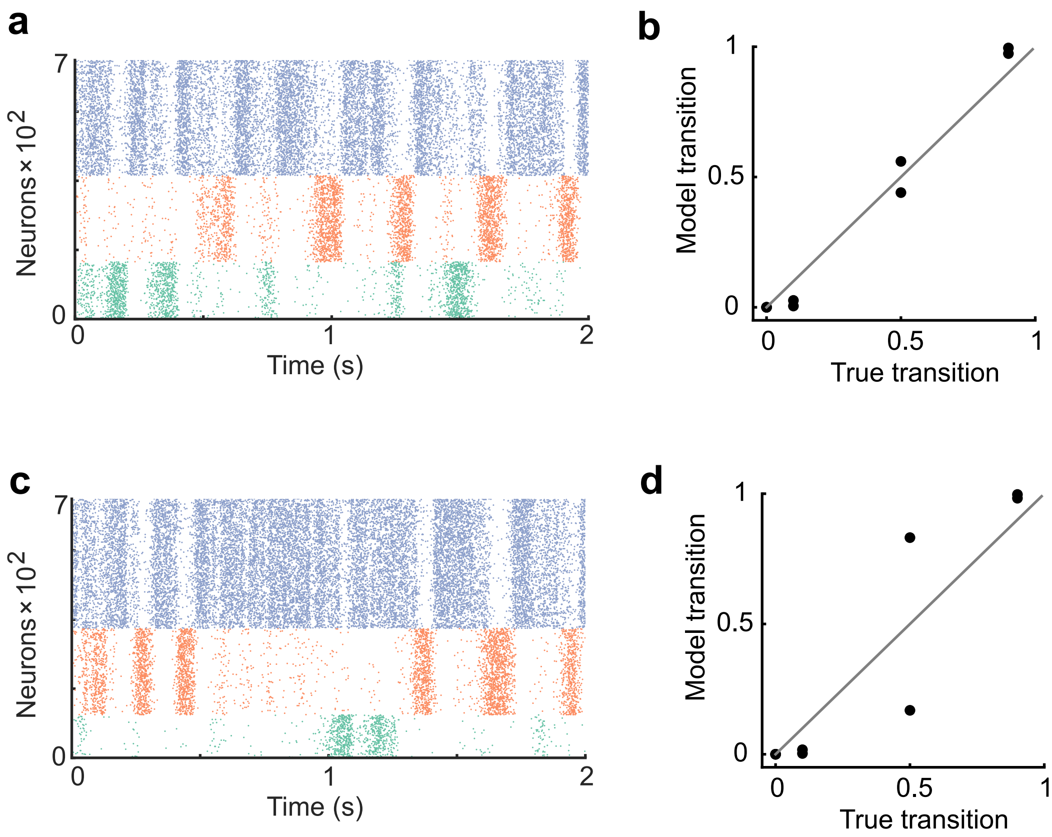


**Supplementary Figure 4. The network performance dependence on cell assembly size.** (a) The network was trained with the assembly size ratio of 1:1.5:2. (b) In the case of a, the spontaneous activity after training reproduced an appropriate transition statistics. (c) The network was trained with the assembly size ratio of 1:2:3. (d) In the case of c, the network showed less performance.
