## Supplementary Figure 5 for "Embedding stochastic dynamics of the environment in spontaneous activity by prediction-based plasticity"

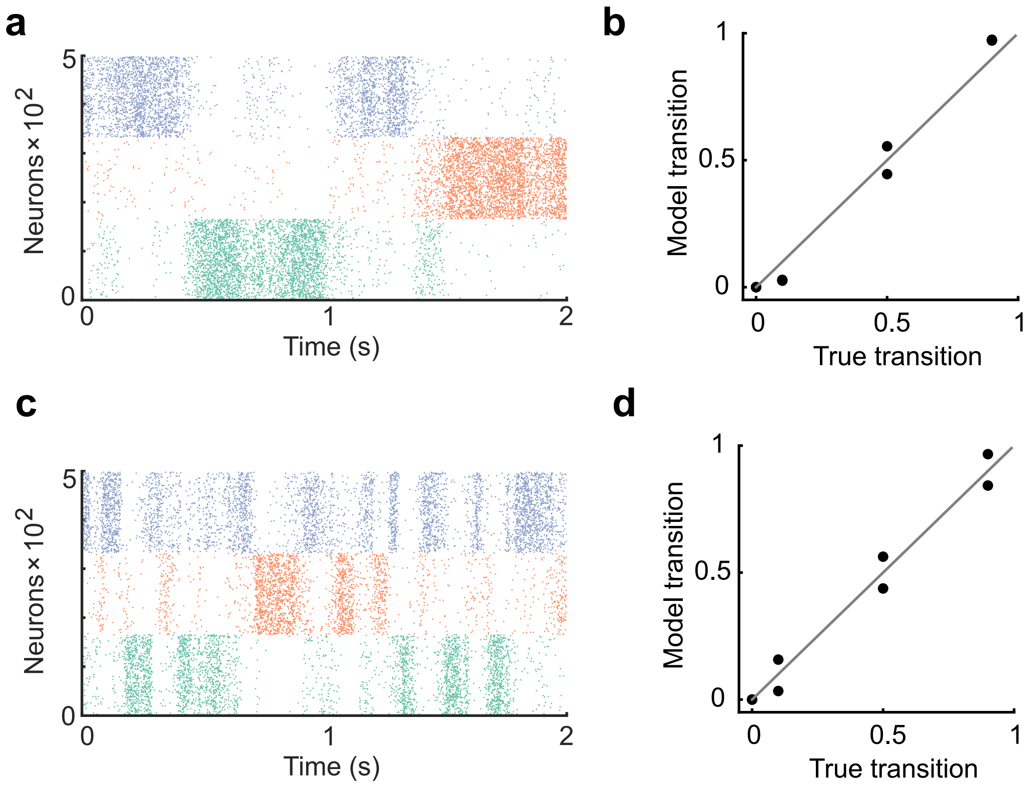


**Supplementary Figure 5. The learning rate controls the duration of cell assembly reactivations.** (a) The learning rate of all plasticity was made half that of the original settings in Figure 2. (b) In the case of a, the spontaneous activity reproduced the transition statistics of the external stimulus patterns. (c) The learning rate of all plasticity was made twice that of the original settings in Figure 2. The duration of assembly reactivations was shorter than in a. (d) Spontaneous activity reproduced transition statistics of external stimulus patterns.
