## Supplementary Figure 6 for "Embedding stochastic dynamics of the environment in spontaneous activity by prediction-based plasticity"

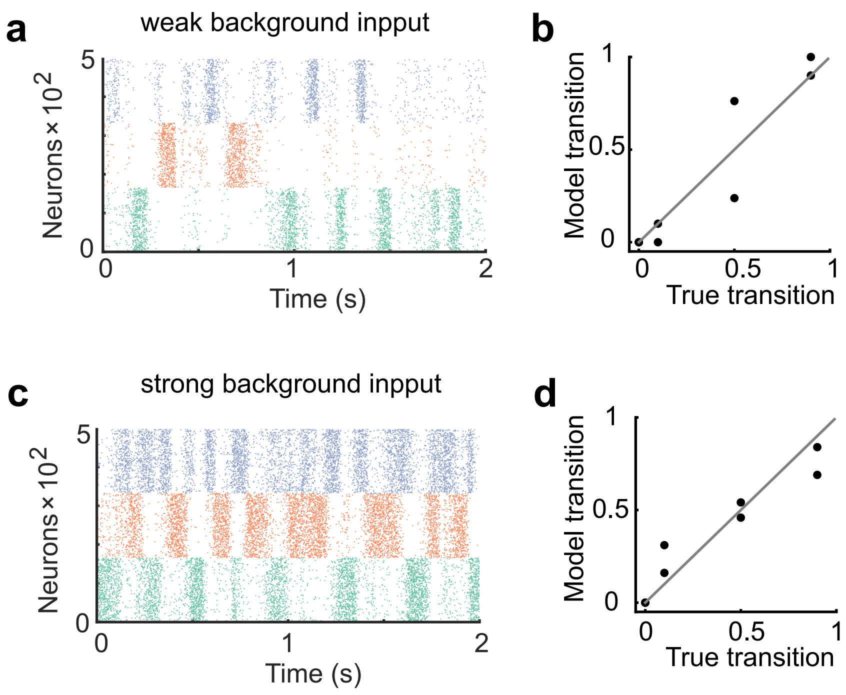


**Supplementary Figure 6. The network performance dependence on the strength of background input.** (a) The network was trained and tested with half the strength of the background input compared to the case in Figure 2. (b) The network showed worse performance than the case shown in Figure 2. (c)The network was trained and tested with double the strength of the background input compared to the case in Figure 2. (d) The network exhibited more uniform transitions compared to the case in Figure 2.
