## Supplementary Figure 7 for "Embedding stochastic dynamics of the environment in spontaneous activity by prediction-based plasticity"

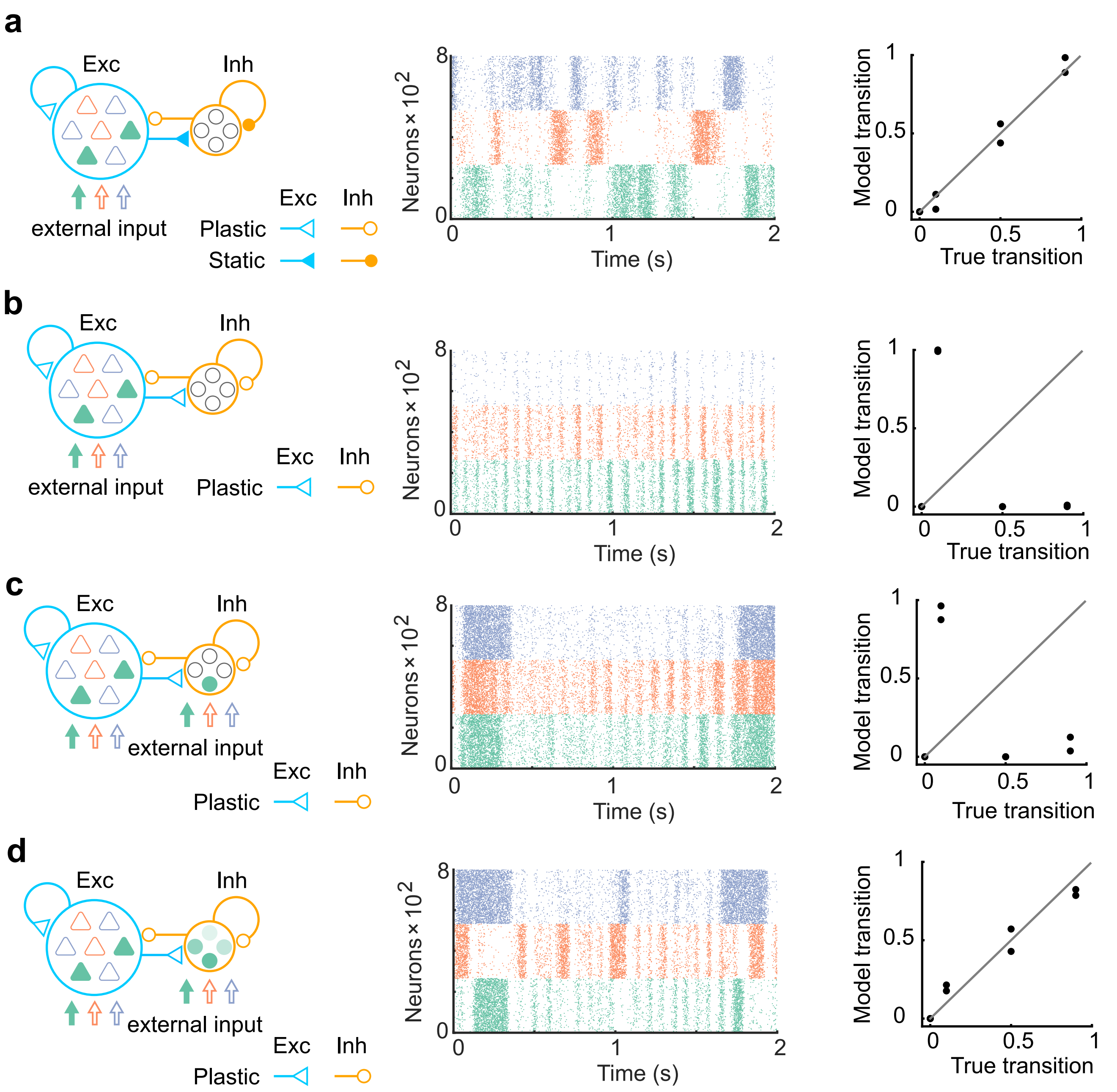


**Supplementary Figure 7. The networks with more biologically plausible architectures.** (a) The network consists of 80% of excitatory neurons and 20% of inhibitory neurons was trained. Same as in Figure1, synapses projecting on excitatory neurons only were trained (left). The network after training showed spontaneous activity with appropriate transition statistics (middle, bottom). (b) Same with a, but all synapses were assumed to be plastic. The network spontaneous activity did not show appropriate transitions. (c) Same with b, but all network neurons receive the external input. The network spontaneous activity did not show appropriate transitions. (d) Same with c, but all inhibitory neurons had mixed selectivity. Here, we assumed that when each state is presented, all inhibitory neurons are driven with a randomly assigned intensity between 0 and 2. The spontaneous activity showed appropriate transitions in this case.
