## Supplementary Figure 8 for "Embedding stochastic dynamics of the environment in spontaneous activity by prediction-based plasticity"

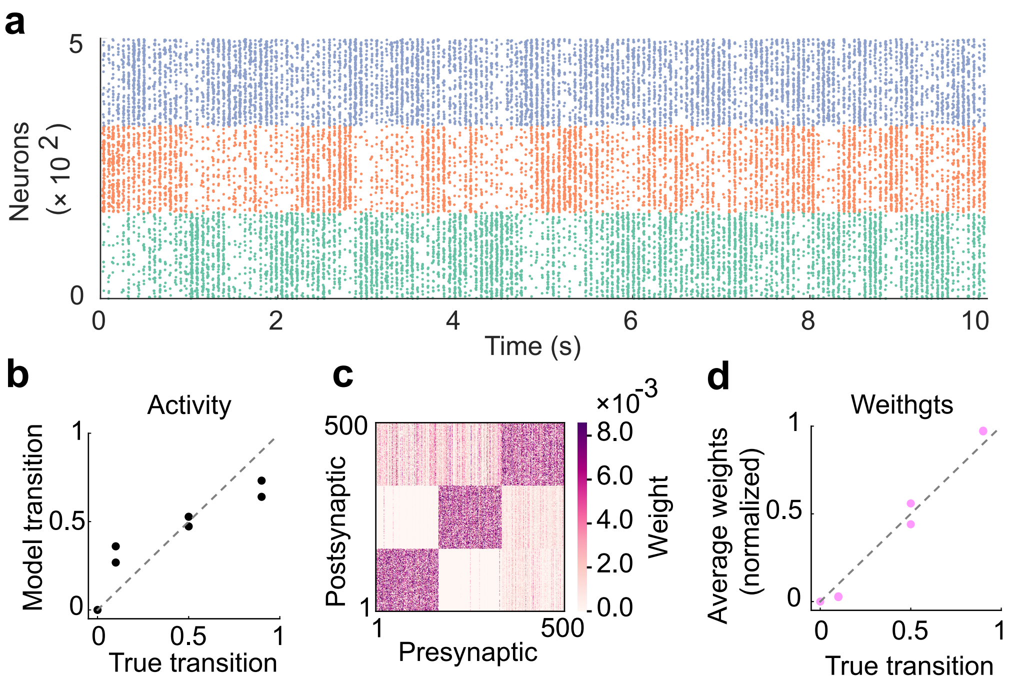


**Supplementary Figure 8. The role of inhibitory plasticity in transition probability learning.** (a) Example spontaneous assembly of a model without inhibitory plasticity is shown. (b) Relationship between the transition statistics of stimulus patterns and that of replayed assemblies. (c) Learned excitatory weights. (d) Relationship between the strength of excitatory synapses between assemblies and true transition probabilities between patterns.
