## Supplementary Figure 9 for "Embedding stochastic dynamics of the environment in spontaneous activity by prediction-based plasticity"

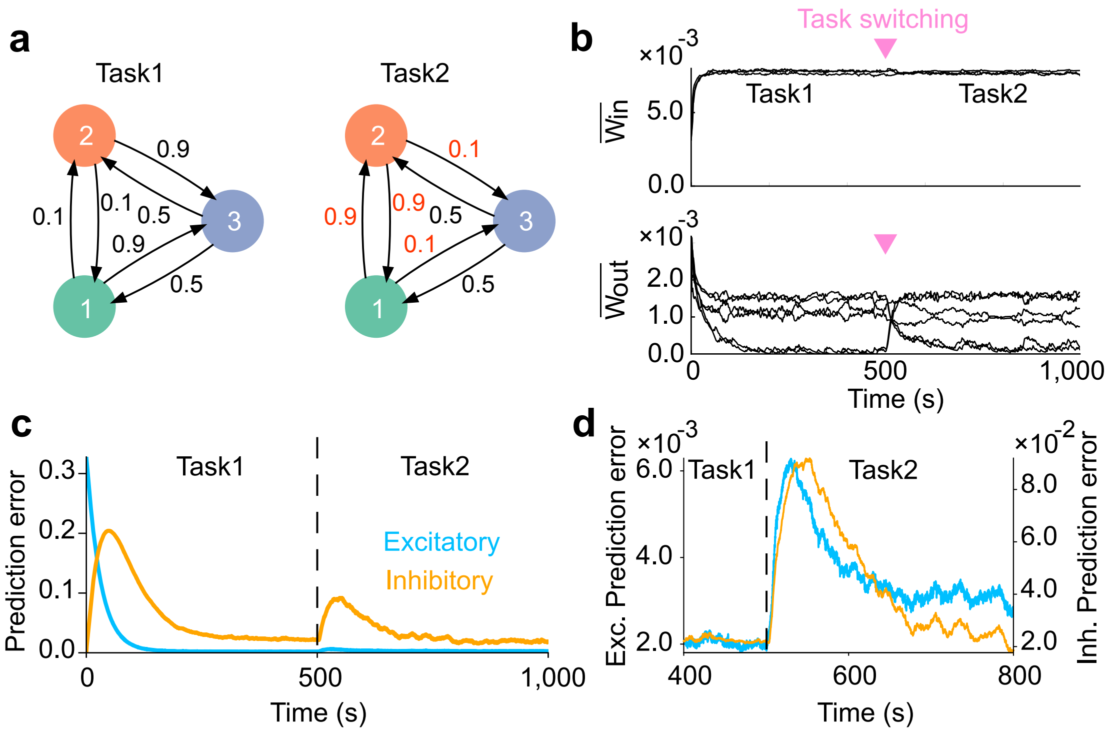


**Supplementary Figure 9. Network adaptation to task switching.** (a) Two types of tasks were considered. (b) Strength of within- (top) and that of between-assembly excitatory synapses (bottom) during learning are shown. Switching from task1 to task2 was occured at the middle of learning phase (inverted triangles). Between-assembly connectivity reorganized once the task switching occurred. (c) Dynamics of prediction error for excitatory (blue) and inhibitory (orange) plasticity are shown. (d) Magnified versions of c are shown. Both errors show an abrupt increase immediately after task switching, followed by a gradual decay. In c and d, errors were calculated as an averages over five independent simulations.
