## Supplementary Figure 10 for "Embedding stochastic dynamics of the environment in spontaneous activity by prediction-based plasticity"

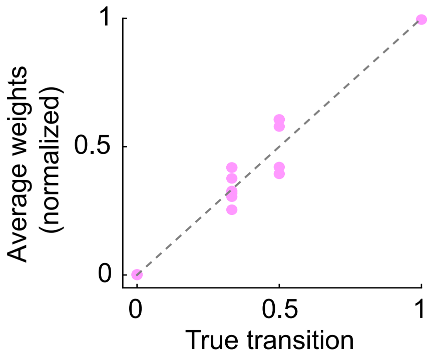


**Supplementary Figure 10. The excitatory synapses learned transition structures of complex task shown in Fig.4a.**
