## Supplementary Figure 11 for "Embedding stochastic dynamics of the environment in spontaneous activity by prediction-based plasticity"

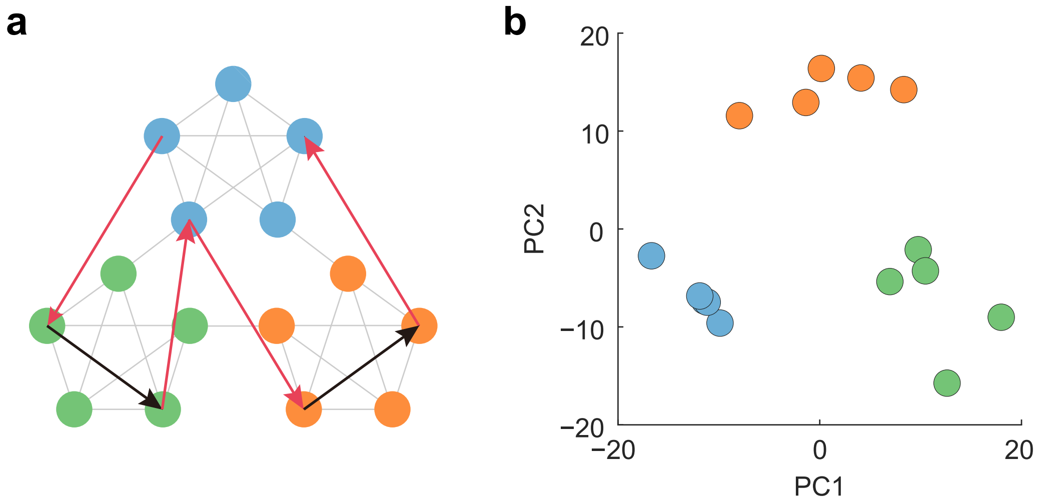


**Supplementary Figure 11. (a) Scrambled structure in which presentation rule of patterns violates temporal community structure. Scrambled sequence consists of both learned (black arrows) and untrained (red arrows) transitions, which viorates the community structure. The network underwent scrambled task only after it learned community structure shown in Fig.4d. (b)Low dimensional representation of activity patterns evoked by scrambled sequence.**
